# Repeatable individual variation in body temperature is associated with cold tolerance and exercise-induced hyperthermia in zebra finches

**DOI:** 10.64898/2026.09.12.750407

**Authors:** Elana Rae Engert, Fredrik Andreasson, Andreas Nord, Jan-Åke Nilsson

## Abstract

In endotherms, body temperature is regulated by balancing the rates of metabolic heat production and dissipation. Birds and mammals maintain high and relatively constant body temperatures, but there is growing evidence that body temperature and heterothermy vary considerably between individuals of a population. Based on the interdependence of metabolic rate and body temperature, variation in body temperature could be expected to be related to metabolic performance. In this study, we aimed to determine if body temperature is a variable and repeatable trait in captive zebra finches (*Taeniopygia guttata*) and if body temperature could be related to different metrics of performance. We measured body temperature for 14 days in in caged male zebra finches to assess within- and among-individual variation and repeatability, as well as during measurements of three metabolic traits – basal (BMR), summit cold-induced (M_sum_) and maximum exercise-induced (MMR) metabolic rate. Median T_b_ (R = 0.48), and peak T_b_ (R = 0.53) body temperature were repeatable and positively correlated, suggesting that the birds show consistent differences in body temperature. We found that birds with higher daytime body temperatures showed higher thermoregulatory performance in a cold challenge and had higher body temperature during forced exercise. However, daytime body temperature was not related to MMR or BMR. Our study is one of few that have measured the repeatability of body temperature in birds. Our results show that individuals may have distinct body temperature phenotypes which could be related to metabolic and thermoregulatory performance. However, additional research is needed to disentangle the interrelated effects of metabolic capacity, body temperature, and heat dissipation on performance and how this might change with thermal conditions and activity levels.

## Introduction

In endotherms, body heat is produced endogenously via metabolism and retained by insulation. Body temperature is regulated through balancing heat production and dissipation, via feedback systems controlled in the hypothalamus that control metabolic heat production, changes to insulation and vascular tone, evaporative heat loss and behavior (Mitchell et al., 2025). Metabolic heat production is the main thermoregulatory mechanism when temperatures decrease, and one of the main determinants of maintaining a stable body temperature. Heat is also produced during activity, which normally results in increased body temperature. Body temperature is tightly linked to organismal function and performance because it influences the rates and of biochemical reactions and stability of macromolecules (Angilletta et al., 2010). Body temperature is also thought to be a strong determinant of metabolic rate among species when controlling for body size, because it determines the rate of biophysiological processes (Brown et al., 2004; Gillooly et al., 2001), resulting in a feedback loop with metabolism (Clarke, 2017). Therefore, metabolic capacity, body temperature, and environmental temperature are inter-related in endotherms and could contribute to organismal performance through different mechanisms and regulatory pathways.

The interdependence of metabolic rate, body mass, and body temperature can lead to complex and contrasting patterns in the scaling of body temperature with other physiological traits in endotherms (Clarke & Rothery, 2008). In line with this, species, populations, and individuals can show substantial variation in their normothermic body temperature and flexibility of body temperature (Angilletta et al., 2010; Boyles et al., 2013; Freeman et al., 2022; McKechnie et al., 2021). In the few studies that have investigated variation in body temperature within and among individuals, these traits have also been found to be individually repeatable in small mammals and birds (Boratyński et al., 2019; Nilsson & Nord, 2018; Tapper et al., 2021). That individuals show consistent differences in body temperature and heterothermy is particularly interesting if these traits are related to metrics of performance, such as the capacity for thermogenesis or exercise, that could presumably impact fitness (Angilletta et al., 2010).

Songbirds maintain relatively high normothermic and maximum body temperatures, capacity for heterothermy, and mass-specific metabolic rates compared to other taxonomic groups (McKechnie et al., 2021). Performance, measured during a cold challenge or during intense exercise, could be hypothesized to be limited by metabolic capacity, heat dissipation capacity, or body temperature in different scenarios. For example, the maximum cold-induced metabolic rate (summit metabolic rate, M_sum_) is the main determinant of body temperature stability during a cold tolerance in small songbirds (Briga & Verhulst 2017, Stager et al 2020) and may improve survival in cold environments (Petit et al., 2017). When the capacity for heat production is exceeded by heat loss, body temperature and metabolic rate steadily decrease, resulting in hypothermia. On the other hand, when heat production via metabolism exceeds the rate of heat dissipation, body temperature generally increases with air temperature, resulting in hyperthermia (McKechnie et al., 2021). Birds routinely become hyperthermic during periods of high activity, such as when provisioning nestlings (Nilsson & Nord, 2018; Tapper et al., 2020). During heat stress, the evaporative cooling capacity, the ratio of evaporative heat loss (EHL) to metabolic heat production (MHP) is the main determinant of heat tolerance in dry conditions (McKechnie et al., 2021). The capacity for exercise and thermogenesis seem to share physiological mechanisms, and have been linked to skeletal muscle mass, lipid catabolism and the circulatory and respiratory systems (Bury et al., 2019; Chappell et al., 1999; Petit & Vézina, 2014; Zhang et al., 2015).

Individual variation in body temperature and capacity for heterothermy may also play a role in thermal tolerance and individual performance. For example, a higher body temperature in a cold environment translates to a larger gradient between body and air temperature, and may require higher thermogenic capacity to maintain body temperature in cold environments (Angilletta et al., 2010). In warm environments, individuals with a higher maximum body temperature may endure higher environmental temperatures in humid air, when evaporative cooling is constrained (Freeman et al., 2022). Variation in normothermic body temperature could also be related to the scope for hyperthermia, depending on whether maximum body temperature is individually variable or physiologically or biochemically constrained. Furthermore, if thermoregulatory or exercise performance is body temperature-dependent according to a thermal performance curve similar to ectotherms, metabolic rate could theoretically be constrained if body temperature deviates from thermal optima for performance (Angilletta et al., 2010).

The sources and significance of metabolic rate and body mass have been studied extensively in endotherms. However, the relevance of normothermic body temperature and flexibility of body temperature for thermoregulatory and exercise performance has received comparatively less attention (Angilletta et al., 2010). In this study, we aimed to describe variation in body temperature between and within individuals, and to determine if body temperature could be related to different metrics of performance. To answer these questions, we continuously measured daytime body temperature in captive zebra finches (*Taeniopygia guttata*), as well as T_b_ during measurements of three metabolic traits – BMR, M_sum_ and MMR. Thermoregulatory performance was measured as M_sum_ and cold endurance, while exercise performance was measured as MMR and exercise endurance.

## Methods

Two weeks before the start of data collection, 18 male zebra finches were transported from an outdoor aviary population at Stensoffa ecological field station to an indoor animal facility at Lund University. All zebra finches were kept in a room with four 1 m3 cages with 4 or 5 individuals in each cage in a room kept at 20 ºC for the duration of the experiment. Only males were used in the experiment to avoid birds going into breeding condition, as all birds were kept in one room and zebra finches breed opportunistically. They were kept on a 12-hour light-dark cycle with lights on and off at 07:00 and 19:00. All birds were implanted with a temperature-sensitive PIT tag (LifeChip BioTherm, Destron Fearing, South St Paul, MN, USA) in the intraperitoneal cavity using the method described in Persson et al. (2024) at least two days before respirometry measurements began. The tags measured 2.12 × 13 mm and weighed 0.26 g.

### Metabolic Measurements

Metabolic measurements were taken from January 14th – 26th 2024. We measured two individuals per day and all three metabolic rates (BMR, M_sum_ and MMR) were measured within 24 hours. BMR was measured overnight, followed by M_sum_ in the morning and MMR in the afternoon. Birds were allowed to rest in cages with *ad libitum* access to food and water for 1 – 3 hours between BMR and M_sum_ measurements, and for 3 – 5 hours between M_sum_ and MMR measurements.

We measured BMR overnight to ensure that birds were in their rest phase and in a post-absorptive state. We took two birds at a time from the animal housing facility to the respirometry lab between 17:00 and 18:00. Each bird was placed in a 1.2 l glass chamber (15 × 15 × 11 cm) with a plastic locking lid which was covered in aluminum tape and spray-painted matte black. The chambers were placed in a dark climate cabinet set to 35 ºC, which is in thermoneutrality for zebra finches (Briga & Verhulst, 2017). Dry atmospheric air (Drierite™; Fisher Scientific, Göteborg, Sweden) was pushed at a mean rate of 854.2 ml m-1 ± SD: 75.0 ml m-1 (standard temperature and pressure, dry, STPD) measured by a FB8 mass flow meter (Sable Systems, Las Vegas, NV, USA). Excurrent air was subsampled at a rate of 165.1 ml m-1 ± SD: 10.2 ml m-1 STPD using a SS-4 sub-sampler (Sable Systems). O2, CO2, WVP (water vapor pressure) in the subsampled air stream and BP (barometric pressure) were measured using an FMS (Field Metabolic System, Sable Systems). Measurements started with a 15-minute measurement of baseline air, followed by a 15-minute measurement of sample air in each respirometry chamber, and a second 15-minute baseline, repeated throughout the night. The following morning at 07:00, the birds were returned to their cages with access to food and water.

Summit metabolic rate (M_sum_) measurements started in the morning between 09:00 and 11:30. M_sum_ was measured in a helox (79% helium and 21% oxygen) environment because thermal conductance is higher in helox than in air (Thomas et al., 1998), allowing for M_sum_ to be reached at higher air temperatures. Helox gas was pushed into the chamber at a mean rate of 1218.1 ± SD: 63.0 ml m-1, measured using an Alicat 0-20 SLPM flowmeter (Alicat Scientific, Tucson, AZ, USA) set to measure the flow rate of the helox gas mixture, and was subsampled at a mean rate of 114.4 ± SD: 46.99 ml m-1. The air temperature inside each chamber was measured continuously using a 36-gauge type T thermocouple connected to a TC-2000 thermocouple box (Sable Systems). Excurrent gas concentrations, BP and air temperature were measured in the same way as in BMR measurements. Baseline air was measured for 5 minutes at the beginning and end of each data recording. Birds were weighed on an electronic scale before and after respirometry measurements. We measured one bird at a time with the same chambers that were used for BMR measurements. After a bird was placed in the chamber, we started a sliding-cold exposure protocol that started at 10 ºC and decreased by 3 ºC every 20 minutes (Swanson et al., 1996). Tests were concluded when metabolic rate decreased steadily and birds became hypothermic, indicating that M_sum_ had been reached. The mean ± SD body temperature of birds when measurements ended was 35.5 ± 0.7 ºC. Birds were then removed from the chamber and placed in another climate chamber set to 25 ºC for about 10 minutes, for them to rewarm to normal body temperatures. They were then returned to their cages for 3 – 5 hours with *ad libitum* access to food and water.

In the afternoon after M_sum_ measurements, we used a hop-flutter wheel to measure MMR (for a description and dimensions, see Engert et al., 2026b). The hop-flutter wheel causes the bird to hop and flap continuously during measurements. The wheel was inside of the climate chamber which was set to 20 ºC. Atmospheric air was dried using Drierite™ and was pushed into the wheel at a mean rate of 2986.7 ± SD: 110.8 ml m-1 STPD and was subsampled at a mean flow rate of 298.6 ± SD: 15.4 ml m-1 STPD. The mass of each bird was measured with a digital scale before and after MMR measurements. After a 15-minute acclimation period, the wheel started to rotate at a speed of 0.4m/s. Every 3 min, the speed was increased by 0.1m/s until it reached a maximum rotation speed of 1.1m/s. The wheel was stopped when the bird started to show signs of exhaustion (sliding inside the wheel or rolling over itself). After stopping the wheel, we continued to measure metabolic rate during a 15-minute recovery period, after which the bird was returned to the animal housing facility. Baseline air was measured for 5 minutes at the start and end of each data recording period. MMR data measured in a different cohort of male zebra finches at 20 ºC as a part of another study (Engert et al., 2026a preprint) were combined with the data from the present study to increase sample size.

### Body temperature

Body temperature was measured continuously during BMR, M_sum_ and MMR measurements using HPR Plus data loggers (Biomark, Boise, ID, USA) with the antennae placed within close range of the respirometry chambers. Daytime body temperature was measured for two weeks in the animal housing facility by placing the antennae on a perch in each cage. The antennae have a range of approximately 10 centimeters, so T_b_ was recorded when birds came within range of the antennae in the cages. Body temperature was variable and peaked periodically in a synchronized way for most individuals each day, even when birds were left undisturbed in the animal housing facility (supplementary material, Figure S1).

### Statistical Analyses

Data analyses were carried out in R (R Core Team, 2022). Respirometry data was recorded and periods of interest were extracted using ExpeData (v1.9.27). Metabolic rate was calculated as the rate of oxygen consumption (VO_2_) and converted to watts as described in (Engert et al., in press). For BMR, the mean of the most level 10-minute period was extracted from each hourly recording, and the minimum value of these selections was considered as BMR. For M_sum_ and MMR, the maximum rolling 5-minute mean was selected out of the entire period of cold exposure or exercise

(Figure 1). Cold endurance and exercise endurance were calculated as the total duration of the cold or exercise challenge (in minutes).

**Figure 1.**
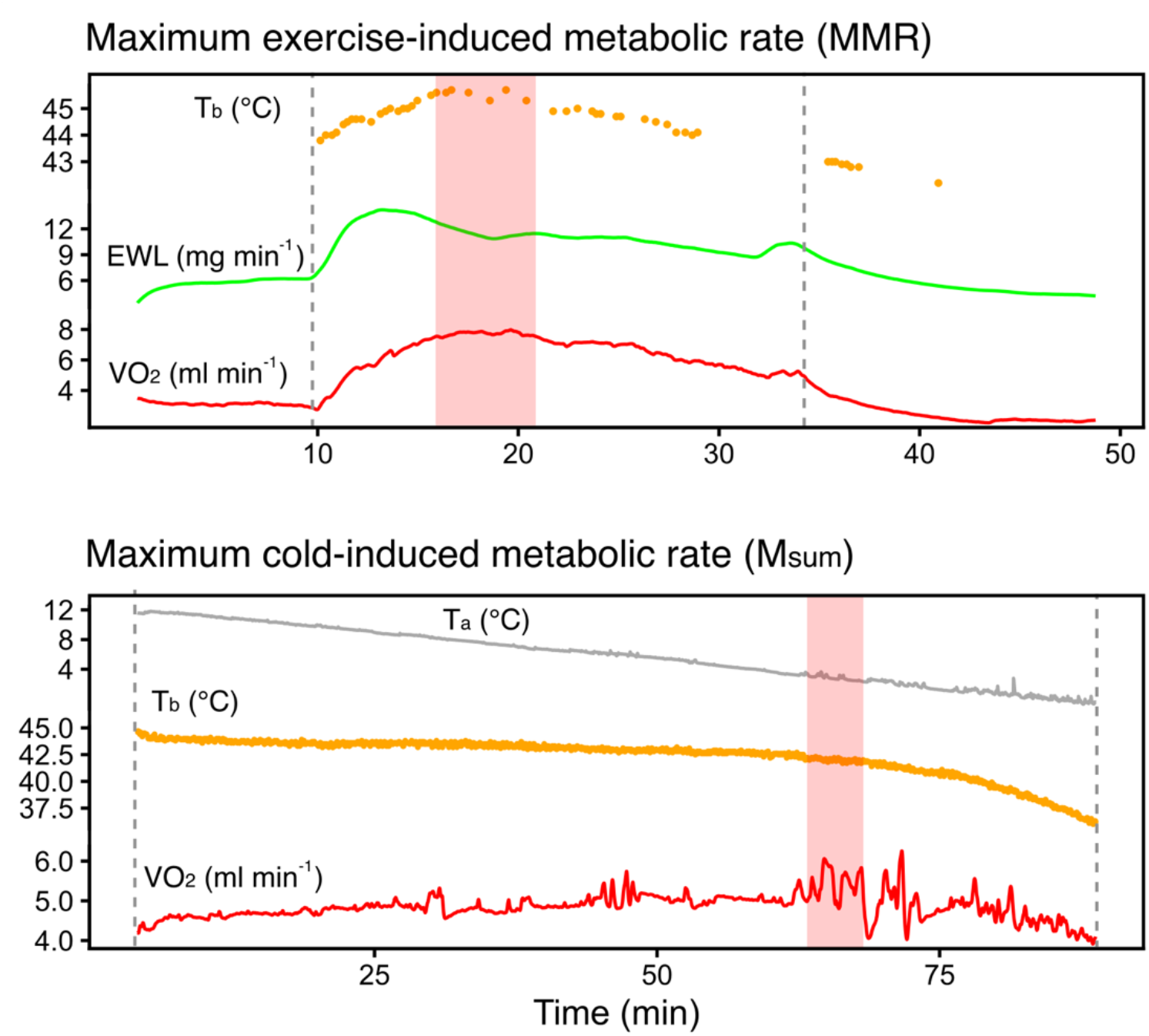
Representative plot of selections of periods of interest during MMR and M_sum_ measurements. The maximum 5-minute rolling mean VO_2_ was selected (selection shown in red shaded areas) from the whole period of exercise or cold exposure (dashed lines). The mean body temperature (orange points) within this selection was calculated as T_b_ during MMR (T_b MMR_) or M_sum_ (T_b Msum_). Evaporative water loss (EWL) is also shown as a green trace during MMR and air temperature (T_a_) in a helox environment is shown as a gray trace during M_sum_.

For body temperature data from the animal housing facility, the individual median and maximum body temperature were recorded per day for repeatability analyses and the mean daily median (median T_b_) and maximum T_b_ (peak T_b_) for each individual was calculated for further analysis. We chose to use the median body temperature instead of modal because daily individual body temperature data was not consistently unimodal. Days for specific individuals with low coverage of T_b_ data (< 100 points) were removed from the analysis. Since median daytime and peak body temperature in the cages turned out to be significantly related, we used only one of these variables in analyses and chose the one that was hypothesized to be of greater relevance for that particular trait. Because birds are not expected to become hyperthermic during BMR and M_sum_, we used median T_b_ in association with these variables. For MMR, we used the peak T_b_ because birds exhibited hyperthermia during exercise.

We investigated relationships between metabolic rates using reduced major axis regressions (RMA) using the *smatr* package (Warton et al., 2012). We used the *rptR* package (Stoffel et al., 2017) to calculate the intra-class correlation coefficient (ICC) of individual median and peak body temperatures. We estimated the individual difference in daily median and peak T_b_ using a linear mixed model with T_b_ as the response variable, the trait as a categorical factor (median or peak T_b_), with individual random intercepts and slopes. A random intercept for day nested within ID was included to account for paired data within days. We tested if random slopes significantly improved the model fit by fitting two models using restricted maximum likelihood (REML), with and without random slopes, which we compared using a likelihood ratio test. We tested for a correlation between median and peak T_b_ using a Pearson correlation. Relationships between T_b_ and metabolic rates measured at the same time were investigated using linear regressions, with T_b_ as the response variable. The relationships between daytime T_b_ and other traits were analyzed as linear regressions or as multiple regressions including body mass.

## Results

### Standardized metabolic rates and body temperature during measurements

All three metabolic rates (BMR, MMR, and M_sum_) were significantly, positively related to one another (Figure 2). When corrected for body mass, only the maximal metabolic rates, MMR and M_sum_, were positively related. MMR was highly repeatable when measured one day apart (R = 0.931, n = 6). While we did not find that BMR was related to body temperature when BMR was measured (T_b BMR_), M_sum_ and MMR were both positively related to T_b_ during when metabolic rate was measured (T_b Msum_ and T_b MMR_, respectively, Table 2). T_b_ was similarly related to metabolic intensity, i.e. body mass-independent residual metabolic rates (supporting information, Table S1).

**Figure 2.**
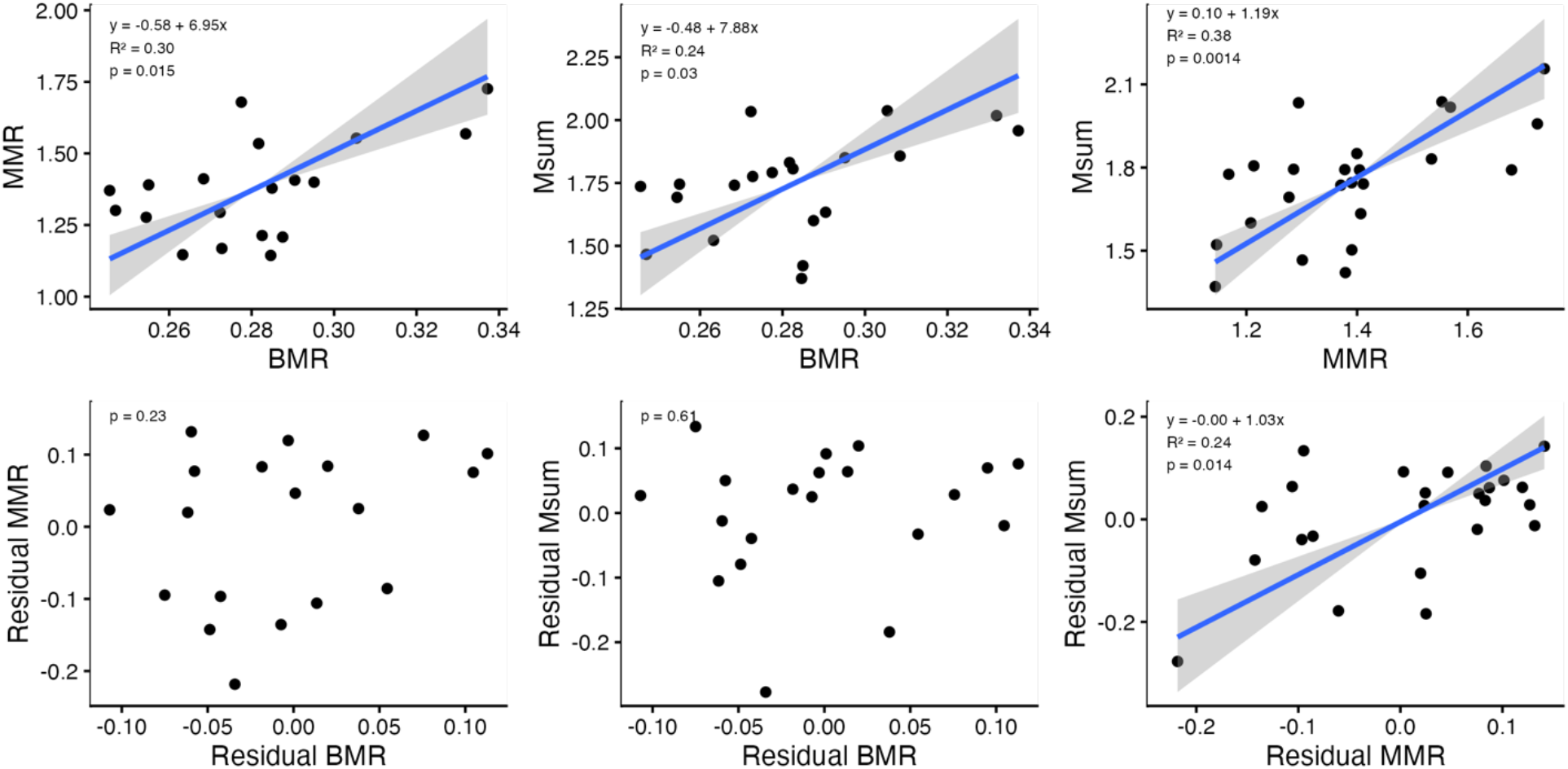
Relationships between whole-animal (top) and body mass-corrected residual (bottom) metabolic rates in captive zebra finches. Estimates, slopes, and p-values are the results of Ranged Major Axis (RMA).

**Table 1.**
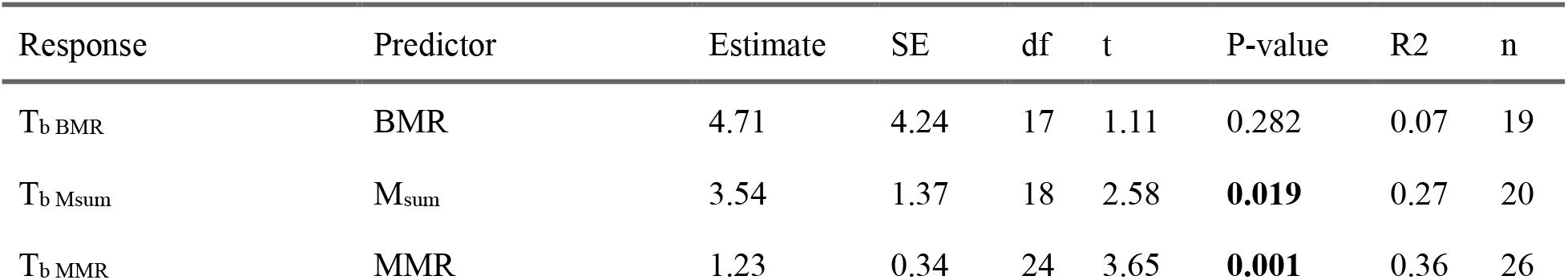
Results of linear models of basal metabolic rate (BMR), summit metabolic rate (M_sum_), and maximum exercise-induced metabolic rate (MMR) with body temperature at the time of measurement and body mass as predictors.

**Table 2.** Results of linear regressions of basal metabolic rate (BMR), summit metabolic rate (M_sum_), and maximum exercise-induced metabolic rate (MMR) and body temperature at the time of metabolic measurements (T_b BMR_, T_b Msum_, and T_b MMR_) with median or peak daytime body temperature as the predictor variable.

| Response | Predictor | Estimate | SE | df | t | P-value | R2 | n |
| --- | --- | --- | --- | --- | --- | --- | --- | --- |
| $T_{b \text{ BMR}}$ | Median daytime $T_b$ | -0.27 | 0.34 | 17 | -0.78 | 0.444 | 0.04 | 19 |
| $T_{b \text{ Msum}}$ | Median daytime $T_b$ | 0.95 | 1.02 | 18 | 0.93 | 0.367 | 0.04 | 20 |
| $T_{b \text{ MMR}}$ | Peak daytime $T_b$ | 0.66 | 0.26 | 12 | 2.50 | **0.028** | 0.34 | 14 |
| BMR | Median daytime $T_b$ | 0.01 | 0.02 | 18 | 0.45 | 0.659 | 0.01 | 20 |
| $M_{\text{sum}}$ | Median daytime $T_b$ | 0.28 | 0.14 | 18 | 2.02 | 0.058 | 0.18 | 20 |
| Cold endurance | Median daytime $T_b$ | 29.11 | 14.88 | 18 | 1.96 | 0.066 | 0.17 | 19 |
| MMR | Peak daytime $T_b$ | 0.09 | 0.08 | 17 | 1.02 | 0.323 | 0.06 | 19 |
| Exercise endurance | Peak daytime $T_b$ | 1.42 | 2.04 | 17 | 0.69 | 0.497 | 0.03 | 19 |

### Body temperature

Daytime T_b_ showed both within- and among-individual variation (Figure 3A). Median T_b_ varied from 42.5 – 43.7 (mean: 43.1) and peak T_b_ ranged from 44.0 – 45.7 (mean: 45.0). Median T_b_ (R = 0.48), and peak T_b_ (R = 0.53) were individually repeatable across days (Figure 3B) and median T_b_ and peak T_b_ were positively correlated (Pearson correlation: r = 0.50, t_18_ = 2.5, p = 0.023, Figure 3C).

**Figure 3.**
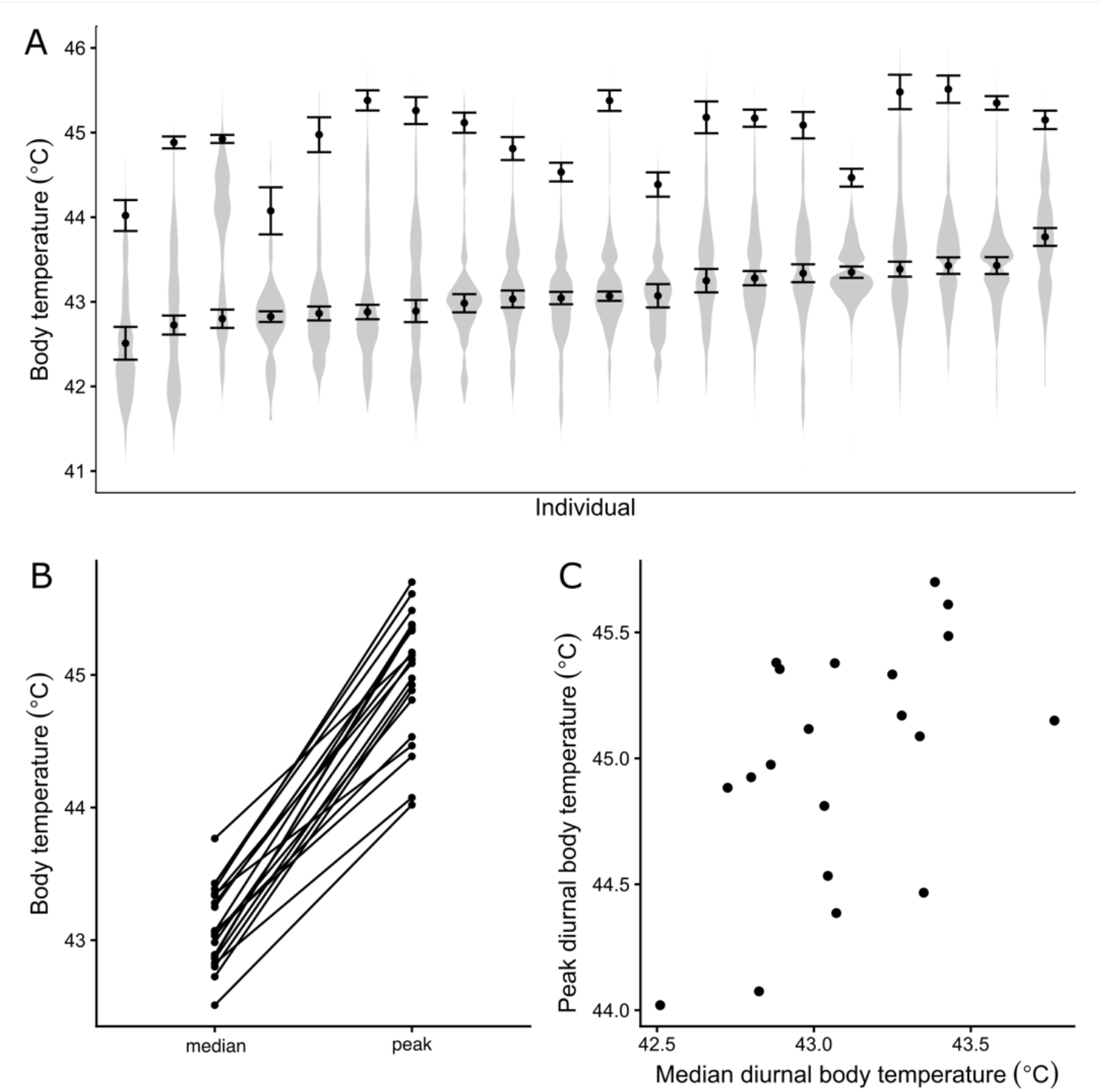
A: Individual variation in daytime core body temperature in captive zebra finches housed in an animal facility at 20 ºC, arranged by median T_b_. Points and error bars show the mean and SE of daily median T_b_ and daily peak T_b_ per individual. B: Median and peak T_b_ were repeatable and individuals showed variation in slope but median T_b_ was not correlated with the slope. C: Median and peak T_b_ were positively correlated.

Peak T_b_ was estimated as being 1.90 ± 0.09 ºC (mean ± SE) higher than median T_b_ (F_1, 18.78_ = 406.23, p < 0.001). There was support for among-individual variation in the magnitude of difference between median and peak T_b_ (SD of random slopes = 0.39 ºC). Inclusion of a random slope term significantly improved the model fit relative to a model including random intercepts only (likelihood-ratio test: χ^2^ = 46.1, p < 0.001). We did not find a correlation between individual intercepts and slopes (r = -0.04), indicating that median T_b_ was not associated with magnitude of difference between median and peak T_b_.

We did not find a relationship between body mass and daytime T_b_ (linear regression: p = 0.30) or Peak T_b_ (linear regression: p = 0.78). T_b BMR_ and T_b Msum_ were not found to be related to median T_b_, but T_b MMR_ increased with peak T_b_ (Table 2). M_sum_ and cold endurance increased with median T_b_, but these effects were marginally significant, and M_sum_ was not significantly related to median T_b_ when variation attributable to body mass was accounted for in the model (Table S2). We did not find a relationship between BMR and median T_b_, or between MMR or exercise endurance and peak T_b_.

## Discussion

Our findings contribute to a small number of studies that have measured repeatability and flexibility of body temperature in birds. We found that caged zebra finches exhibited high within-individual variation in body temperature with regular, synchronized peaks that often reached the presumed maximum T_b_ (44 – 45 ºC, Pessato et al., 2022; Wojciechowski et al., 2021). Moreover, that daytime T_b_ and peak T_b_, were repeatable and positively correlated suggests that individuals with relatively high median T_b_ throughout the day also had a relatively high peak T_b_ which could point to distinct T_b_ phenotypes. We also found that while individuals varied in their body temperature flexibility, a high median T_b_ did not seem to constrain the scope for hyperthermia. The results align with evidence showing that individual tree swallows (*Tachycineta bicolor*) consistently differed in body temperature across air temperatures, with males showing individual variation in slopes (Tapper et al., 2021). MMR was also highly repeatable (R = 0.931) when measured one day apart in this study. This implies that MMR is a reliable measure of exercise performance and adds to evidence that metabolic rates are individually consistent traits (reviewed in McKechnie & Swanson, 2010). MMR has also been found to be repeatable in free-living birds across seasons (Engert et al., in press) but not within a breeding season (Engert et al., 2026b). However, both metabolic rates (Swanson, 2010) and thermal sensitivity (Angilletta et al., 2010) are expected to be subject to acclimatization, which could affect repeatability in natural settings.

We found that minimal and maximal measures of metabolic rates were positively related to each other (BMR – M_sum_, BMR – MMR), but residual metabolic rates were not related after accounting for differences in body mass. This is similar to what others have found in passerine birds (Engert et al., in press; Swanson et al., 2012). This suggests that minimal and maximal metabolic rates show positive co-variation at the organismal level but this can also be explained by body mass. However, the two maximal metabolic rates (MMR – M_sum_) were positively related both at the organismal level and relative to body mass. This aligns with evidence that M_sum_ and MMR are functionally related (Petit & Vézina, 2014). BMR is likely related to the mass of digestive organs and related to overall food intake while M_sum_ can vary independently of mass due to changes in metabolic intensity in the muscles (Barceló et al., 2017) which, by association, could be true of MMR as well.

We found evidence that individual variation in T_b_ could be related to metabolic capacity in the context of a cold challenge. Daytime T_b_ was marginally positively related to M_sum_ and cold endurance. M_sum_ was also positively related to T_b Msum_. Thus, birds with a higher daytime T_b_ were more cold-tolerant and maintained a higher T_b_ during a cold-challenge. This could indicate that higher daytime body temperature improves the capacity for thermogenesis, or birds with higher thermogenic capacity are also warmer throughout the day, even when they aren’t cold stressed. An alternative explanation is that those individuals require a higher thermogenic capacity to maintain their body temperature at a higher level when cold-stressed due to a larger gradient between air temperature and body temperature.

Contrary to our expectations, we did not find a relationship between daytime T_b_ and T_b_ during BMR. T_b_ and metabolic rate generally decrease during the rest phase, but our results suggest that individual T_b_ is not consistent between day and night. We also did not find a relationship between BMR or mass-independent residual BMR and T_b_ when they were measured at the same time, so this was likely due to differences in insulation (Scholander et al., 1950). The lack of a pattern could also be explained if there is lower thermoregulatory precision in metabolic rate or passive heat loss when resting in the thermoneutral zone, compared to when cold- or heat-challenged or during activity. This shows that BMR and T_b_ measured during rest phase in thermoneutrality may not be reliably predicted by daytime T_b_.

Birds with higher peak T_b_ during the day sustained higher T_b_ during forced exercise (T_b MMR_, Table 2) and MMR was positively related to T_b_ when MMR was measured. This is perhaps unsurprising, because heat produced during activity cannot always be dissipated fast enough, which leads to exercise-induced hyperthermia (Speakman & Król, 2010). A higher peak T_b_ did not seem to confer higher performance in this study, however, as peak T_b_ was not related to MMR itself. When comparing individuals, it can be difficult to tease apart how metabolic rate, T_b_, and exercise performance are related, due to possible differences in aerobic capacity, capacity to dissipate heat, and thermal tolerance between individuals.

Based on the overall pattern, one possible interpretation is that hyperthermia constrains metabolic rate. However, air temperature could affect both MMR and T_b_ directly or indirectly, as birds could also respond to increased air temperatures by reducing metabolic rate before T_b_ increases to the point where it would affect performance (Tapper et al., 2021). However, peak T_b_ measured in caged birds was higher than T_b_ measured during MMR, which suggests that birds were not constrained by the upper limits to body temperature during exercise (in 20 ºC) in this study (Figure S2). In a previous study of MMR in zebra finches where T_a_ was sequentially increased, body temperature increased and MMR and metabolic scope decreased with increasing T_a_ (Engert et al., 2026a preprint). Furthermore, MMR (r = 0.59) and T_b_ (r = 0.43) were repeatable, meaning that individuals varied in MMR and T_b_ but responded consistently to increased T_a_ relative to one another (Engert et al., 2026a preprint). This also suggests that individuals vary in their heat dissipation rate, for example due to differences in body size, insulation, evaporative cooling capacity, or area of thermal windows.

## Conclusions

In our study, we found support for our hypothesis that T_b_ is a repeatable and could be related to performance when subjected to a cold-challenge or hyperthermia during forced exercise. Thus, our findings strengthen support for consistent individual physiological syndromes (Ricklefs & Wikelski, 2002), but further research is needed to determine if there are ecological implications associated with differences in both median body temperature and exercise-induced hyperthermia. Metabolic rates and daytime body temperatures also seemed to vary independently of one another, with the exception of M_sum_, for which 18% of variation could be explained by variation in median T_b_ in this study.

When body temperature is maintained within safe bounds, we presume that performance is largely determined by metabolic capacity, which results in a decoupling between T_a_ and T_b_ in endotherms. However, when body temperature rises or falls to the extreme ends of thermal tolerance, endotherms are expected to respond with decreased performance, similarly to ectotherms (Angilletta et al., 2010; James & Tallis, 2019; Levesque & Marshall, 2021). During activity in the heat, however, metabolic heat production can be modulated as a thermoregulatory mechanism before T_b_ rises to a point that would constrain performance (Fuller et al., 1998). Thus, T_a_ could affect metabolic rate and T_b_ simultaneously, and performance could also be affected by evaporative cooling capacity and insulation, making it difficult to disentangle the limits to exercise capacity (Levesque & Marshall, 2021). To determine if T_b_ directly affects performance during exercise, future studies could focus on manipulating T_b_ independently of T_a_ to disentangle the inter-related effects of T_a_, T_b_, and metabolic performance.

## Acknowledgements

This study was supported by the Swedish Research Council (Vetenskapsrådet) to J-ÅN (2021-05467) and AN (2020-04686). ERE was supported by Stiftelsen Lars Hiertas Minne (FO2022-0336), Lunds Djurskyddsfond (50/22, 80/23, 63/24) and the Royal Physiographic Society of Lund (2023-44251).

## Data availability statement

All data and script for replication of the results, tables and figures will be made publicly available upon acceptance of this manuscript for publication in a scientific journal.

## References

Angilletta, M. J., Cooper, B. S., Schuler, M. S., & Boyles, J. G. (2010). The evolution of thermal physiology in endotherms. Frontiers in Bioscience E, 2, 861–881. 10.2741/e148

Barceló, G., Love, O. P., & Vézina, F. (2017). Uncoupling Basal and Summit Metabolic Rates in White-Throated Sparrows: Digestive Demand Drives Maintenance Costs, but Changes in Muscle Mass Are Not Needed to Improve Thermogenic Capacity. Physiological and Biochemical Zoology, 90(2), 153–165. 10.1086/689290

Boratyński, J. S., Iwińska, K., & Bogdanowicz, W. (2019). An intra-population heterothermy continuum: notable repeatability of body temperature variation in food-deprived yellow-necked mice. Journal of Experimental Biology, 222(6), jeb197152. 10.1242/jeb.197152

Boyles, J. G., Thompson, A. B., McKechnie, A. E., Malan, E., Humphries, M. M., & Careau, V. (2013). A global heterothermic continuum in mammals. Global Ecology and Biogeography, 22(9), 1029–1039. 10.1111/geb.12077

Briga, M., & Verhulst, S. (2017). Individual variation in metabolic reaction norms over ambient temperature causes low correlation between basal and standard metabolic rate. Journal of Experimental Biology, 220(18), 3280–3289. 10.1242/jeb.160069

Brown, J. H., Gillooly, J. F., Allen, A. P., Savage, V. M., & West, G. B. (2004). Toward a metabolic theory of ecology. Ecology, 85(7), 1771–1789. 10.1890/03-9000

Bury, A., Niedojadlo, J., Sadowska, E. T., Bauchinger, U., & Cichon, M. (2019). Contrasting response of haematological variables between long-term training and short exercise bouts in zebra finches (Taeniopygia guttata). Journal of Experimental Biology, 222(4), Article jeb193227. 10.1242/jeb.193227

Chappell, M. A., Bech, C., & Buttemer, W. A. (1999). The relationship of central and peripheral organ masses to aerobic performance variation in house sparrows. Journal of Experimental Biology, 202(17), 2269–2279. 10.1242/jeb.202.17.2269

Clarke, A. (2017). Principles of Thermal Ecology: Temperature, Energy and Life. Oxford University Press. 10.1093/oso/9780199551668.001.0001

Clarke, A., & Rothery, P. (2008). Scaling of body temperature in mammals and birds. Functional Ecology, 22(1), 58–67. 10.1111/j.1365-2435.2007.01341.x

Engert, E. R., Grange, M., Nord, A., Andreasson, F., & Nilsson, J.-Å. (in press). Fitness and flexibility of metabolic phenotypes in a temperate resident bird. Functional Ecology.

Engert, E. R., Nilsson, A., Andreasson, F., Nord, A., & Nilsson, J.-A. (2026a). Air temperature limits aerobic capacity and scope in active birds. bioRxiv, 2026.2009.2009.750338. 10.64898/2026.09.09.750338

Engert, E. R., Nord, A., Andreasson, F., & Nilsson, J.-Å. (2026b). Flexibility of exercise capacity during nestling feeding in blue tits. Journal of Experimental Biology, 229(7), jeb251043. 10.1242/jeb.251043

Freeman, M. T., Czenze, Z. J., Schoeman, K., & McKechnie, A. E. (2022). Adaptive variation in the upper limits of avian body temperature. Proceedings of the National Academy of Sciences, 119(26), e2116645119. 10.1073/pnas.2116645119

Fuller, A., Carter, R. N., & Mitchell, D. (1998). Brain and abdominal temperatures at fatigue in rats exercising in the heat. Journal of Applied Physiology, 84(3), 877–883. 10.1152/jappl.1998.84.3.877

Gillooly, J. F., Brown, J. H., West, G. B., Savage, V. M., & Charnov, E. L. (2001). Effects of Size and Temperature on Metabolic Rate. Science, 293(5538), 2248–2251. 10.1126/science.1061967

James, R. S., & Tallis, J. (2019). The likely effects of thermal climate change on vertebrate skeletal muscle mechanics with possible consequences for animal movement and behaviour. Conservation Physiology, 7(1), coz066. 10.1093/conphys/coz066

Levesque, D. L., & Marshall, K. E. (2021). Do endotherms have thermal performance curves? Journal of Experimental Biology, 224(3). 10.1242/jeb.141309

McKechnie, A. E., Gerson, A. R., & Wolf, B. O. (2021). Thermoregulation in desert birds: scaling and phylogenetic variation in heat tolerance and evaporative cooling. Journal of Experimental Biology, 224, jeb229211. 10.1242/jeb.229211

McKechnie, A. E., & Swanson, D. L. (2010). Sources and significance of variation in basal, summit and maximal metabolic rates in birds. Current Zoology, 56(6), 741–758. 10.1093/czoolo/56.6.741

Mitchell, D., Fuller, A., Snelling, E. P., Tattersall, G. J., Hetem, R. S., & Maloney, S. K. (2025). Revisiting concepts of thermal physiology: understanding negative feedback and set-point in mammals, birds, and lizards. Biological Reviews, n/a(n/a). 10.1111/brv.70002

Nilsson, J.-Å., & Nord, A. (2018). Testing the heat dissipation limit theory in a breeding passerine. Proceedings of the Royal Society B: Biological Sciences, 285(1878), 20180652. 10.1098/rspb.2018.0652

Persson, E., Cuív, C. O., & Nord, A. (2024). Thermoregulatory consequences of growing up during a heatwave or a cold snap in Japanese quail. Journal of Experimental Biology, 227(2), Article jeb246876. 10.1242/jeb.246876

Pessato, A., McKechnie, A. E., & Mariette, M. M. (2022). A prenatal acoustic signal of heat affects thermoregulation capacities at adulthood in an arid-adapted bird. Scientific Reports, 12(1). 10.1038/s41598-022-09761-1

Petit, M., Clavijo-Baquet, S., & Vézina, F. (2017). Increasing Winter Maximal Metabolic Rate Improves Intrawinter Survival in Small Birds. Physiological and Biochemical Zoology, 90(2), 166–177. 10.1086/689274

Petit, M., & Vézina, F. (2014). Phenotype manipulations confirm the role of pectoral muscles and haematocrit in avian maximal thermogenic capacity. Journal of Experimental Biology, 217(6), 824–830. 10.1242/jeb.095703

R Core Team. (2022). R: A language and environment for statistical computing. In R Foundation for Statistical Computing. https://www.R-project.org/

Ricklefs, R. E., & Wikelski, M. (2002). The physiology/life-history nexus. Trends in Ecology & Evolution, 17(10), 462–468. 10.1016/s0169-5347(02)02578-8

Scholander, P. F., Walters, V., Hock, R., & Irving, L. (1950). Body Insulation of Some Arctic and Tropical Mammals and Birds. Biological Bulletin, 99(2), 225–236. 10.2307/1538740

Speakman, J. R., & Król, E. (2010). Maximal heat dissipation capacity and hyperthermia risk: neglected key factors in the ecology of endotherms. Journal of Animal Ecology, 79(4), 726–746. 10.1111/j.1365-2656.2010.01689.x

Stoffel, M. A., Nakagawa, S., & Schielzeth, H. (2017). rptR: Repeatability estimation and variance decomposition by generalized linear mixed-effects models. Methods in Ecology and Evolution, 8(11), 1639–1644. 10.1111/2041-210X.12797

Swanson, D. L. (2010). Seasonal Metabolic Variation in Birds: Functional and Mechanistic Correlates. In C. F. Thompson (Ed.), Current Ornithology Volume 17 (pp. 75–129). Springer New York. 10.1007/978-1-4419-6421-2_3

Swanson, D. L., Drymalski, M. W., & Brown, J. R. (1996). Sliding vs static cold exposure and the measurement of summit metabolism in birds. Journal of Thermal Biology, 21(4), 221–226. 10.1016/0306-4565(96)00005-8

Swanson, D. L., Thomas, N. E., Liknes, E. T., & Cooper, S. J. (2012). Intraspecific Correlations of Basal and Maximal Metabolic Rates in Birds and the Aerobic Capacity Model for the Evolution of Endothermy. PLOS ONE, 7(3), e34271. 10.1371/journal.pone.0034271

Tapper, S., Nocera, J. J., & Burness, G. (2020). Experimental evidence that hyperthermia limits offspring provisioning in a temperate-breeding bird. Royal Society Open Science, 7(10). 10.1098/rsos.201589

Tapper, S., Nocera, J. J., & Burness, G. (2021). Body temperature is a repeatable trait in a free-ranging passerine bird. Journal of Experimental Biology, 224(20), jeb243057. 10.1242/jeb.243057

Thomas, D. W., Andreina Pacheco, M., Fournier, F., & Fortin, D. (1998). Validation of the effect of helox on thermal conductance in homeotherms using heated models. Journal of Thermal Biology, 23(6), 377–380. 10.1016/S0306-4565(98)00028-X

Warton, D. I., Duursma, R. A., Falster, D. S., & Taskinen, S. (2012). smatr 3 -an R package for estimation and inference about allometric lines. Methods in Ecology and Evolution, 3, 257–259.

Wojciechowski, M. S., Kowalczewska, A., Colominas-Ciuró, R., & Jefimow, M. (2021). Phenotypic flexibility in heat production and heat loss in response to thermal and hydric acclimation in the zebra finch, a small arid-zone passerine. Journal of Comparative Physiology B, 191(1), 225–239. 10.1007/s00360-020-01322-0

Zhang, Y., Eyster, K., Liu, J.-S., & Swanson, D. L. (2015). Cross-training in birds: cold and exercise training produce similar changes in maximal metabolic output, muscle masses and myostatin expression in house sparrows (Passer domesticus). The Journal of Experimental Biology, 218(14), 2190–2200. doi:10.1242/jeb.121822

