## supporting information for "Repeatable individual variation in body temperature is associated with cold tolerance and exercise-induced hyperthermia in zebra finches"

### Body temperature is repeatable and related to cold tolerance and exercise-induced hyperthermia in a passerine bird

**Supplementary tables**

Table S1. Results of linear models of mass-independent residual basal metabolic rate (BMR), summit metabolic rate ($M_{sum}$), and maximum exercise-induced metabolic rate (MMR) in zebra finches with body temperature at the time of measurement and body mass as predictors.

| Response | Predictor | Estimate | SE | DF | t | P-value | R2 | n |
|---|---|---|---|---|---|---|---|---|
| $T_{b\ BMR}$ | Residual BMR | 2.34 | 1.57 | 17.00 | 1.49 | 0.155 | 0.12 | 19.00 |
| $T_{b\ Msum}$ | Residual $M_{sum}$ | 7.39 | 2.61 | 18.00 | 2.83 | 0.011 | 0.31 | 20.00 |
| $T_{b\ MMR}$ | Residual MMR | 2.77 | 0.79 | 24.00 | 3.51 | 0.002 | 0.34 | 26.00 |

Table S2. Results of linear models of basal metabolic rate (BMR), summit metabolic rate ($M_{sum}$), and maximum exercise-induced metabolic rate (MMR) and body temperature at the time of metabolic measurements ($T_{b\ BMR}$, $T_{b\ Msum}$, and $T_{b\ MMR}$) in zebra finches with median or peak daytime body temperature and body mass as covariates.

| Predictor | Estimate | SE | t | P-value |
|---|---|---|---|---|
| BMR | | | | |
| daytime $T_b$ | -0.01 | 0.01 | -0.37 | 0.713 |
| body mass | 0.01 | 0.00 | 3.90 | 0.001 |
| $M_{sum}$ | | | | |
| daytime $T_b$ | 0.21 | 0.13 | 1.62 | 0.124 |
| body mass | 0.06 | 0.02 | 2.35 | 0.031 |
| MMR | | | | |
| daytime $T_b$ | 0.07 | | 0.96 | 0.349 |
| body mass | 0.05 | 0.02 | 2.38 | 0.030 |

#### Supplementary figures

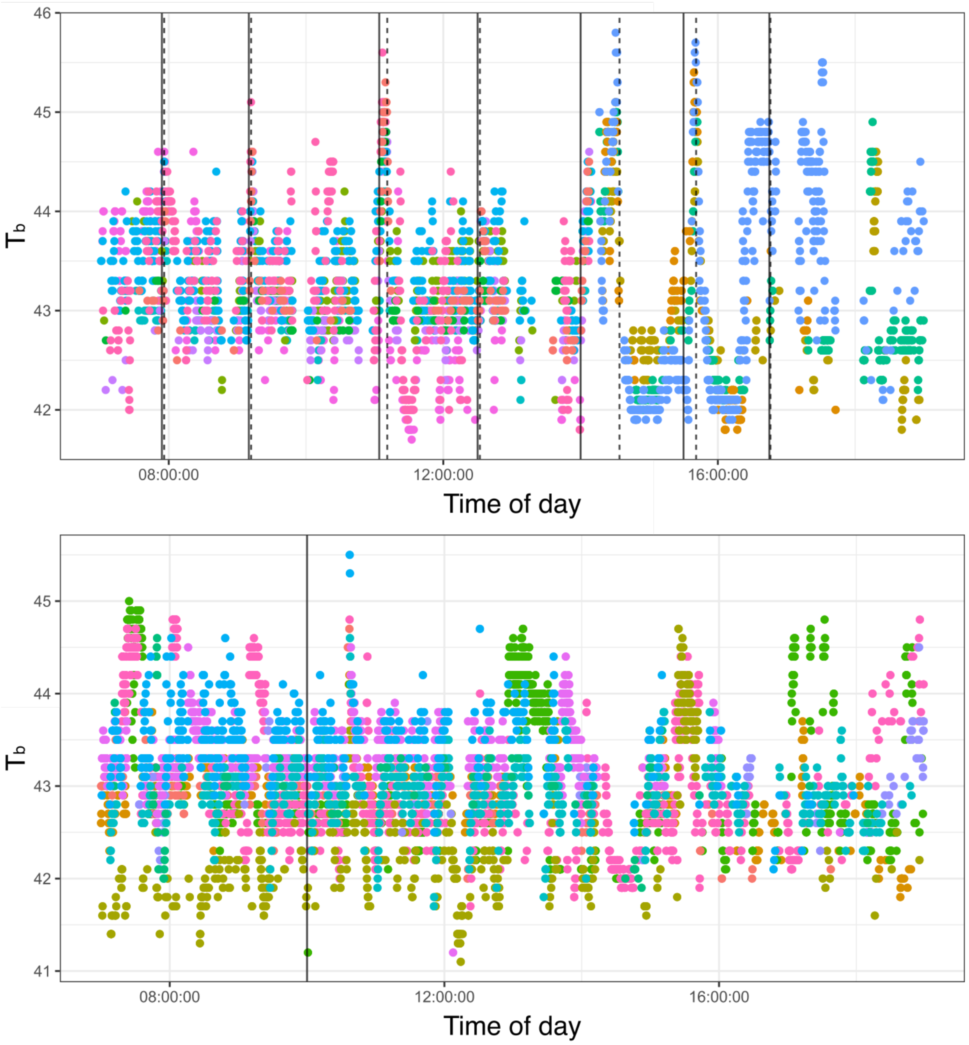


Figure S1. Representative plots of daily body temperature in the animal housing facility showing body temperature of different individual zebra finches (represented by different colors) over time and entry (solid lines) and exit (dashed lines) times of researchers.

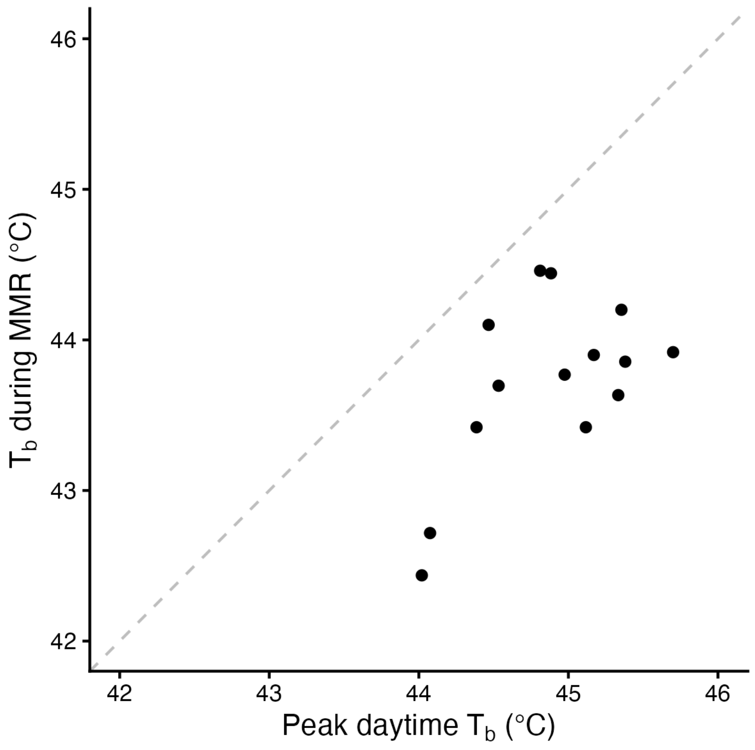


Figure S2. $T_b$ during exercise (MMR) increased with peak daytime body temperature in captive zebra finches, but was lower than during exercise (dashed line is 1:1 reference).
